# Multiple Loci Selection with Multi-way Epistasis in Coalescence with Recombination

**DOI:** 10.1101/2021.02.26.433089

**Authors:** Aritra Bose, Filippo Utro, Daniel E. Platt, Laxmi Parida

## Abstract

As studies move into deeper characterization of the impact of selection through non-neutral mutations in whole genome population genetics, modeling for selection becomes crucial. Moreover, epistasis has long been recognized as a significant component in understanding evolution of complex genetic systems. We present a backward coalescent model EpiSimRA, that builds multiple loci selection, with multi-way (*k*-way) epistasis for any arbitrary *k*. Starting from arbitrary extant populations with epistatic sites, we trace the Ancestral Recombination Graph (ARG), sampling relevant recombination and coalescent events. Our framework allows for studying different complex evolutionary scenarios in the presence of selective sweeps, positive and negative selection with multiway epistasis. We also present a forward counterpart of the coalescent model based on a Wright-Fisher (WF) process which we use as a validation framework, comparing the hallmarks of the ARG between the two. We provide the first framework that allows a nose-to-nose comparison of multiway epistasis in a coalescent simulator with its forward counterpart with respect to the hallmarks of the ARG. We demonstrate through extensive experiments, that EpiSimRA is consistently superior in term of performance (seconds vs. hours) in comparison to the forward model without compromising on its accuracy. EpiSimRA (both backward and forward) source, executable, user manuals are available at: https://github.com/ComputationalGenomics/SimRA.

## 1 Introduction

*Nothing in Biology Makes Sense Except in the Light of Evolution*^1^ and simulating the evolution process, whether of multi-cellular humans, unicellular micro-organisms or even cancer-tumor, continues to be an important device in understanding the observed molecular profiles of populations. These diversity profiles are captured by the genetic variability due to mutations and the change in frequency of alleles within populations over time. The selectively neutral infinite-sites model [18] is often used to analyze this variation [13]. Simulating random populations plays a significant role to investigate effects of complex evolutionary processes on genetic diversity [2]. There are mainly two types of simulation algorithms: backward-time or coalescent and forward-time. The coalescent simulation [19] allow fast approximation of the neutral Wright-Fisher (WF) model with natural selection, shaping patterns of variation in populations. The ARG [11] is a variant of Kingman’s coalescent and is used to reconstruct the *most recent common ancestor* (MRCA), backwards starting from the extant populations or leaves, using coalescent and recombination events. Once it finds the MRCA or if it involves all the trees, the *grand most recent common ancestor* (GMRCA), it traverses the ARG to the extant populations and introduce mutations or other genetic information in the genealogy. Forward-time simulators are more flexible than their backward (coalescent) counterparts in modeling selection along with other factors as it starts from an initial population and tracks its evolution under the influence of various factors such as recombinations, mutations, varying effective population size, fitness effects, etc. It progresses over multiple sequential generations, usually drawing random samples from the last generation to construct an ARG and its hallmarks. Although, coalescent processes are much faster than forward-time simulation algorithms [4].

The classical approach for coalescent simulation as defined by Hudson in the seminal *ms* tool, applied the effects of recombination and coalescence to the ancestors of the samples going back in time in the extant population. This was later computed more efficiently in *msprime* [15] which used a new encoding for correlated trees resulting from simulations of the coalescent with recombination. Some approximations to the coalescent algorithms which are fast also exist such as *SMC* [20], *MaCS* [5], *fastsimcoal* [10]. Many programs were developed to simulate scenarios not captured by ms such as selection [8,25,26,29], demographic inference [8,9] and admixture [3] among others. Coalescent models tracking genealogies in presence of selection can also build an Ancestral Selection Graph (ASG), which is a branching-coalescing random graph within which the genealogy of a sample is embedded [23] conditional on the frequencies of the selected allele of the sample [28]. However, none of these methods take into account epistasis, which has long been recognized as a significant component in understanding genealogies and evolution of complex genetic systems [1]. Here, we present the first coalescent simulator EpiSimRA which captures multiway epistasis i.e. allowing for interaction between alleles in multiple loci under selection. EpiSimRA tracks the ARG from randomly sampled extant populations and unlike ASG not conditional on allele frequencies. It constructs the genealogy dependent on the time to the closest recombination and coalescent event going backwards. Along with this, we also present an alternative, simple forward-time algorithm fwd-EpiSimRA which efficiently simulates epistatic scenarios, to provide a validation framework to the coalescent simulator.

Forward simulators usually track the complete ancestral information, i.e. studying all the lineages that survived until the last generation as a result of recombination events. Only a few forward simulators exist to provide a framework to model epistasis, such as *SELAM* [6], allowing for pairwise epistatic selection to model the process and consequences of admixture or *SLiM* [22,12], which constructs ecologically realistic scenarios while accounting for a host of complex biological processes beyond the WF framework. Specifically its functionality of tree-sequence recording draws parallels to fwd-EpiSimRA in a WF framework, providing support for epistatic interactions. However, none of these packages [6,12] can be used to compare with EpiSimRA as it is not possible to reconstruct the ARG from random extant samples. As fwd-EpiSimRA traces the ARG to obtain the MRCA of the random extant samples and record its hallmarks, we use it for a nose-to-nose comparison with the coalescent simulator as a validation framework. In the remainder of the paper we introduce the coalescent simulator and explain how it tracks multiway epistasis in presence of recombination, followed by an overview of the forward simulator and the ARG tracking algorithm. Thereafter, we show the concordance between the coalescent and forward models for complex evolutionary scenarios and finally concluding with the importance of studying multiway epistasis in simulating real world scenarios of admixture, cryptic relatedness and viral phylodynamics.

## 2 The Coalescent Simulator

The algorithm works back-in-time starting from the present (time 0), moving back into the past. Here we focus on how EpiSimRA is able to simulate multiway epistasis (the interested reader is referred to [3] for the neutral scenario). Let the number of loci under selection be l, possibly with multiway epistasis. As an illustration let *l* be 3 with selection values *s*_1_, *s*_2_ and *s*_3_. The algorithm will assign three random locations on the genetic segment, unless the locations are explicitly specified and we assume that one of the alleles (either major or minor) is under selection while the other is neutral. The possible multiway epistasis are *e*_12_, *e*_13_, *e*_23_ and *e*_123_. If no value is specified then the epistasis is assumed to be neutral. Given this, we get 2^*l*^ possible types of lineages, each of them denoted as *l*_*z*_. Let *l*_0_ be the lineage type with no selection. For the running example, the other lineage types are *l*_1_, *l*_2_, *l*_3_, *l*_12_, *l*_13_, *l*_23_ and *l*_123_. For two lineage types *z*_*a*_ and *z*_*b*_, let

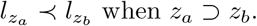

For example, *l*_12_ ≺ *l*_1_ and *l*_12_ ≺ *l*_2_. Also, *l*_123_ ≺ *l*_12_. For the lineage type *z*, let *N*_*z*_ be the effective population size.

### 2.1 Selection Scenarios

The fitness 1 + *s* is the ratio of the probabilities that the selected allele produces an offspring to the neutral allele, which relates to the proportions in generation *t*+1 given proportion in generation *t*. Let *N*_*s*_ be the partially effective population size with the allele under selection and 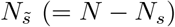 is the partially effective population size with the reference or ancestral allele which is not under selection, giving:

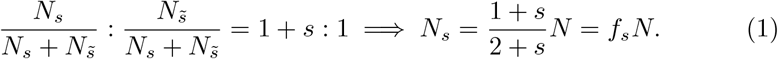

Thus −1 *< s*, extendable to multiple loci with or without epistasis and the fitness defined as

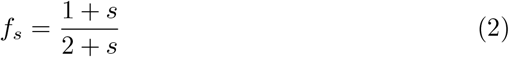

The fitness coefficient is a representative average of the allele frequency of the selected alleles in a generation, *p*. With the allele frequency, the effective population size with selection at a single locus would be 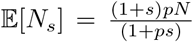 where the fitness 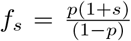. The effective population size is defined as *N*_*z*_ = 2*Nf*′ where *f*′ is the fitness for *l*_*z*_-coalescence in the coalescent simulator and *N*_*z*_ is the effective population size for *l*_*z*_ lineage coalescence with alleles under selection.

The *f*′ varies with neutral or epistatic scenarios for the loci under selection. For a neutral scenario with no selection *f*′ is defined as follows

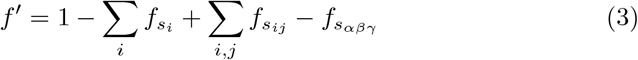

where we remove the fitness effects of odd sites under selection and add the effects from even sites in a simulation scenario with three SNPs (*α, β* and *γ*) are considered to be under selection. Alternatively, for a single locus (*α*) under selection with no epistasis in effect *f*′ will be defined as follows with the signs reversed for even and odd sites under selection

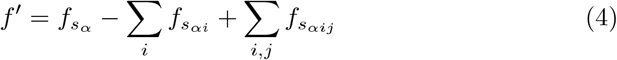

For two or multiple loci under selection there can be two cases with differing *f*_*s*_. We define it as follows:

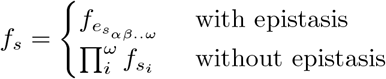

where *e*_*s*_ is the user defined epistatic coefficient when epistasis is in effect across all *ω* sites. For a scenario with all three sites are under selection 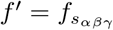.

### 2.2 EpiSimRA: Multiple Loci Selection & Multiway Epistasis

If *s*_*i*_ and *s*_*j*_ are two locations with the minimum (or derived) allele under selection at locus *i* and *j* respectively, then *e*_*ij*_ denotes the epistasis between the two. If it is not explicitly specified then a neutral case (without epistasis) is assumed. The algorithm randomly chooses the location of the SNPs on the genetic segment being simulated.

We assume that no more than one event, coalescent or recombination, occurs at a generation and there is no back mutation, i.e. a base undergoes no more than one mutation in the entire ARG. The mutation and recombination rates are uniform over the segment being simulated. If there is recombination, the lineages are randomly assigned but if *r* = 0, the lineages are so assigned that no pair of types of lineages straddle (either they are disjoint or one is contained in the other). Lineage *l*_0_ corresponds to lineage with no alleles under selection. For each lineage *l*_*z*_, the algorithm only appends each node to a list when a recombination occurs with time *t* > *T*_*z*_ where *T*_*z*_ keeps track of the time to GMRCA. The recombination rate for 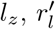 is defined as

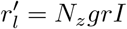

For nodes which are not leaf nodes the length of the genetic material, *s* is not proportional to the recombination rate as the rate is governed by the effective population size *N*_*z*_. The stochastic nature of the method allows for a *Pooled* loop which pools lineages together at each iteration in the minimum of *t*_*z*_, over all lineages *l*_*z*_. For each lineage *t*_*z*_ = *N*_*z*_ ×*t* is computed where *t* is the time to next event using

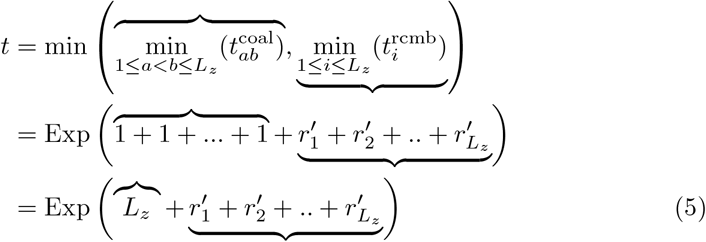

*t*^coal^ is the time to coalescence and similarly *t*^rcmb^ is the time to the next recombination event. Equation 5 computes the minimum time to the next event (coalescent or recombination). When there is only one lineage in the pool, only recombination event can occur.

#### Coalescence event

In a coalescence event *L*_*z*_ is decremented by 1 as two random lineages of type *l*_*z*_ are coalesced into one at time *T*_*z*_ and the outgoing edge of the coalesced node is labeled by lineage *l*_*z*_. If |*L*| = 1, *z* is a singleton label (such as *s*_1_ but not *s*_1_*s*_2_ or *s*_1_*s*_2_*s*_3_), and, there exist no active lineage 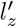 such that *z*′ ≺*z*, then the mutation(s) corresponding to lineage *l*_*z*_ is assigned to this edge (using an approach in [3]) and the label of the outgoing edge of the new node is changed to *l*_0_ and *L* is incremented by 1. Next, *L* is set to 0 and thus the lineage *l*_*z*_ is made inactive.

##### Algorithm 1: EpiSimRA

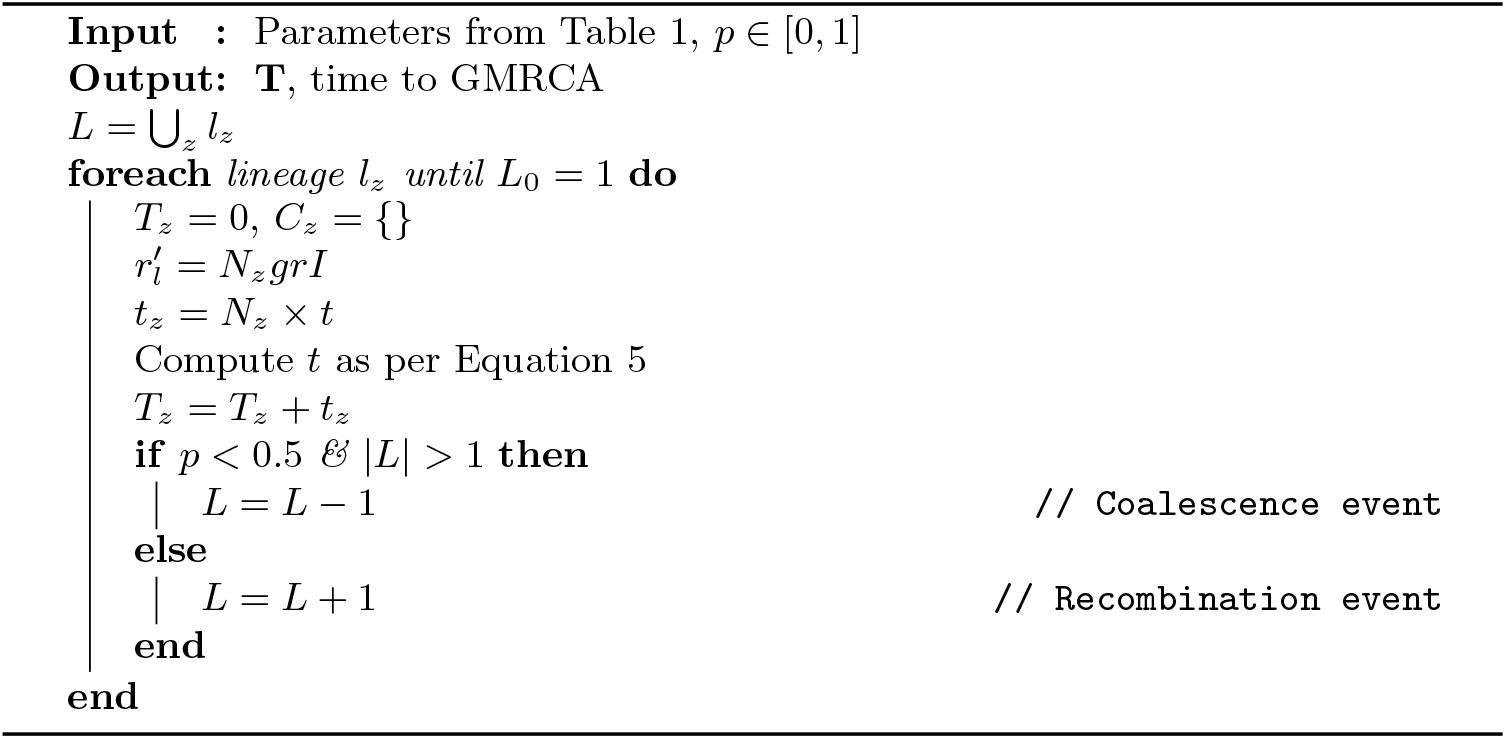

#### Recombination event

In a recombination event a lineage of type *l*_*z*_ is randomly picked and a node *v* is created at *T*_*z*_. The label of *z* is randomly split it into two lineage labels that is compatible with the location of the SNPs on the genetic segment *I* carried by the node *v*. Thereafter, *L* is incremented by 1.

The algorithm for EpiSimRA is described in Algorithm 2.2 and see Appendix for an illustrative example of the algorithm for a three-way epistatic scenario.

## 3 The Forward Simulator

The model simulates evolution for a full population, forward in time with each generation containing *N* equal number of males and females, each carrying two chromosomes (see Appendix for a detailed discussion and extension to selection on multiple loci). The complex evolutionary relationships between generations yields a number of mutations, recombinations, selected allele inheritance, linkage disequilibrium, etc. along the length of chromosome for each individual. This data is recorded in a data structure, which we call the “book of populations”. We trace the lineage of each site along the chromosome while tracing the ‘book’ and constructing the ARG. Inheritance follows the convention of a standard WF model applied to diploid organisms [13], with children randomly picking their parents corresponding to the fitness coefficients when selection is in effect.

### 3.1 Simulating the “Book of Populations”

Each chromosome is represented by the alleles at each locus *l* ∈ [1, *g*], which is randomly assigned initially. We use same notations as defined in section 1 to describe fwd-EpiSimRA. The model assumes that each locus *l* has a fitness function *s*_*l*_(*a*) ∈ ℝ, where *a* is an allele comprising the genotype. An individual *i* with allele *a*_*il*_ at locus *l* is assigned a selection coefficient *s*_*il*_ = *s*(*a*_*il*_) which is user-defined, similar to EpiSimRA. The function *s*(.) denotes the selective pressure and can be varied by intentional specification of recessive, dominant, additive, and other configurations, including homozygous advantage. This function encompasses selection at both single and multiple loci allowing flexible user-defined variations. When selection is not present, we set *s*_*il*_ = 0.

For an individual *i*, the probability that it has children is given by

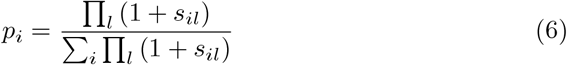

(see Appendix for derivation). In each new generation, as in the WF model, the *N* children pick their parents with replacement according to the parent probabilities *p*_*i*_. The simulation is run for *t* = {0, 1, …, *G*} discrete generations with the *t* = 0 being the base generation, outlined in Figure 1(a).

**Fig. 1.**
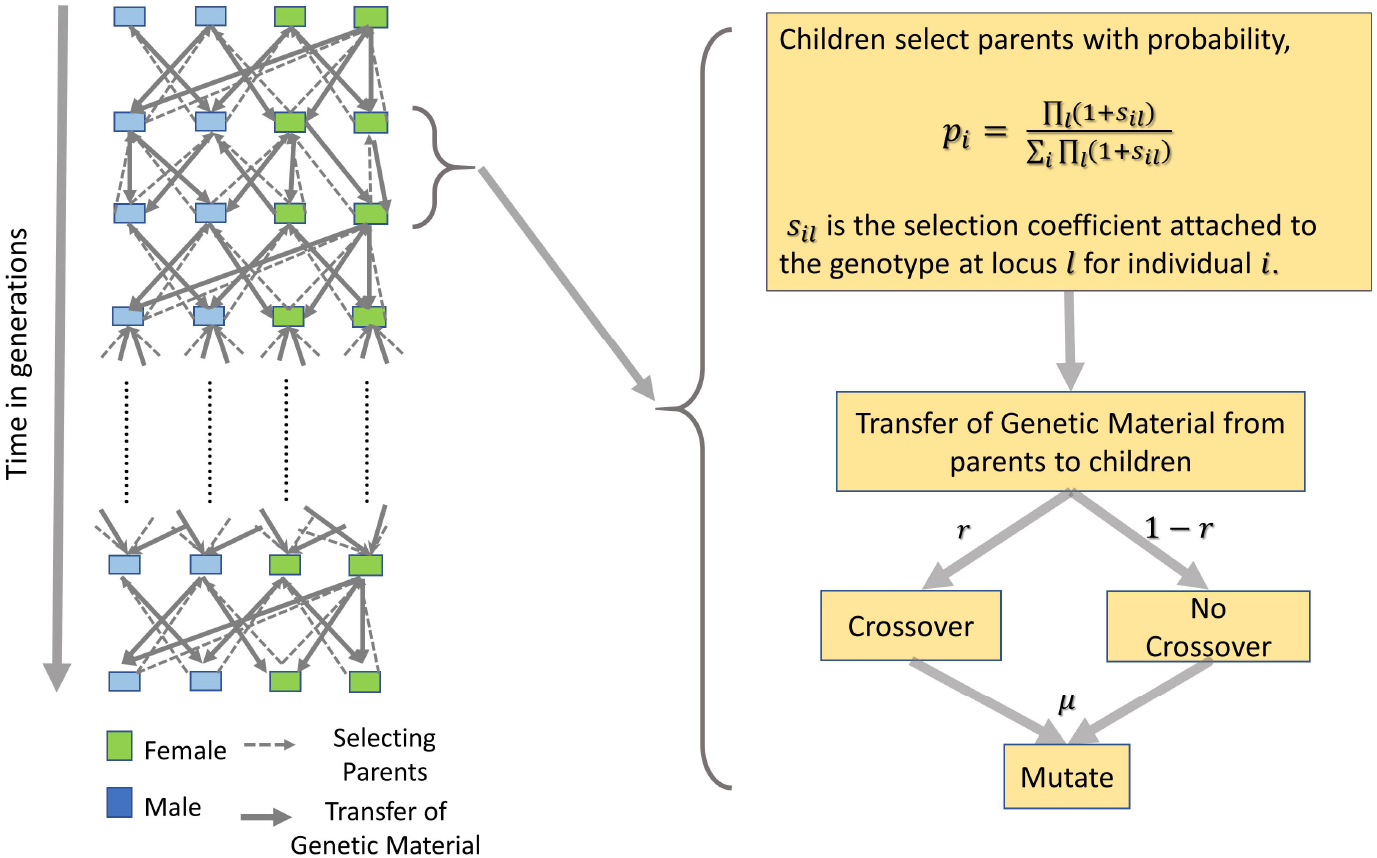
Schematic diagram for simulating the “book of populations” which closely resembles the biological process of evolution.

In each new generation, as in the WF model, the *N* children pick their parents with replacement according to the parent probabilities *p*_*i*_. The simulation is run for *t* ={0, 1, …, *G*} discrete generations with the *t* = 0 being the base generation, outlined in Figure 1(a).

### 3.2 Modeling Multiway Epistasis

Multiway epistasis requires multiple interacting loci with similar selection effects. We assign selection coefficients to interacting sites for *k*-way epistasis, where *k* is the maximum number of interacting sites. Let there be *q* groups of loci, each containing at most *k* elements and we re-compute equation (6) accounting for fitness related to interacting sites as,

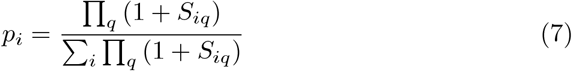

If a group only has one element, that is if the selected locus is non-interacting, then we allow *S* = *s*, the user defined selection coefficient. For all other cases, we select *S* from a matrix or tensor of all possible allele combinations with respect to the number of interacting sites. *S*, the combined fitness coefficient is calculated by taking the fitness product of each interacting site as,

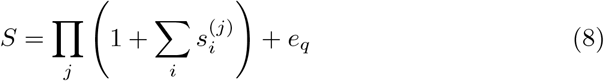

*e*_*q*_ is the epistatic interaction coefficient for each combination of interacting sites and 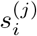 is the selection coefficient at allele *j* in individual *i*’s chromosome.

### 3.3 Tracing the ARG

Detecting the past recombination events from extant sequences and specifying the place of each recombination is well studied [14,21,24]. The ARGs define a genealogical graph for all of the chromosomes in a population. Recent advances in population genetics simulators have resulted in tree-sequence recordings which obtains the genealogical history of all genomes in a simulated population [16]. However, no natural ARG is recorded for the interacting loci with epistasis in effect and randomly sampling populations from extant generation, in forward simulators. It is traced from the “book of populations” from a number of extant haplotypes. We start from *m* randomly selected extant populations and trace the recombination and coalescent events back each generation. We keep a track of each lineage corresponding to every site along the chromosome and stop when we have found a convergence for all lineages. This final coalescent event along the entire “book” is known as MRCA and we output the corresponding ARG.

## 4 Empirical Evaluation

Selection in a diploid heterozygous sample can boost, for one generation, the non-selected chromosome. This can entangle the impact of selection on lineages in the diploid forward model, but not the haploid. We expected the impact of boosted preference to be minimal along any given lineage, since such a boost only occurs for dominant or additive alleles, and then for only one generation. In combinations in a population over time, this effect could be more significant and thus, we sought to test this.

### 4.1 Comparison Study

Comparing the two models under selection calls for an assessment of the values. In both the models, common phenomena such as faster coalescence, decreasing diversity, decreasing number of recombination events occur when we study the individuals under selection. Hence, we compare the *H*, the height of the ARG or the time to MRCA, as it is the most significant hallmark of the common history of a sample. We run simulations for different parameter set-ups for the forward and backward model by running each experiment 100 times. We demonstrate the accuracy of the two algorithms by comparing *H* under different simulation scenarios allowing at most three interacting loci. The simplest scenario in this case is when there is no selection in effect i.e., the neutral coalescent model and selection at a single locus. We show that the two proposed models EpiSimRA and fwd-EpiSimRA show agreement in all of the different epistatic scenarios including selection in single locus.

The results for the complex scenario in this setting, accounting for epistasis with three loci are shown in Fig. 2, where we show the concordance for the forward and backward simulation with box-whisker plots, QQ-plots, CDF plots and PP plots (Figure B7 in Appendix). To obtain further validation we observed similar agreement in Kolmogorov-Smirnov (KS) test on the distributions of *H* as returned by fwd-EpiSimRA and EpiSimRA for all scenarios. We found that for each, the null hypothesis that the two samples are drawn from the same distribution is never rejected and the test statistic is very small (Table B2 in Appendix).

**Fig. 2.**
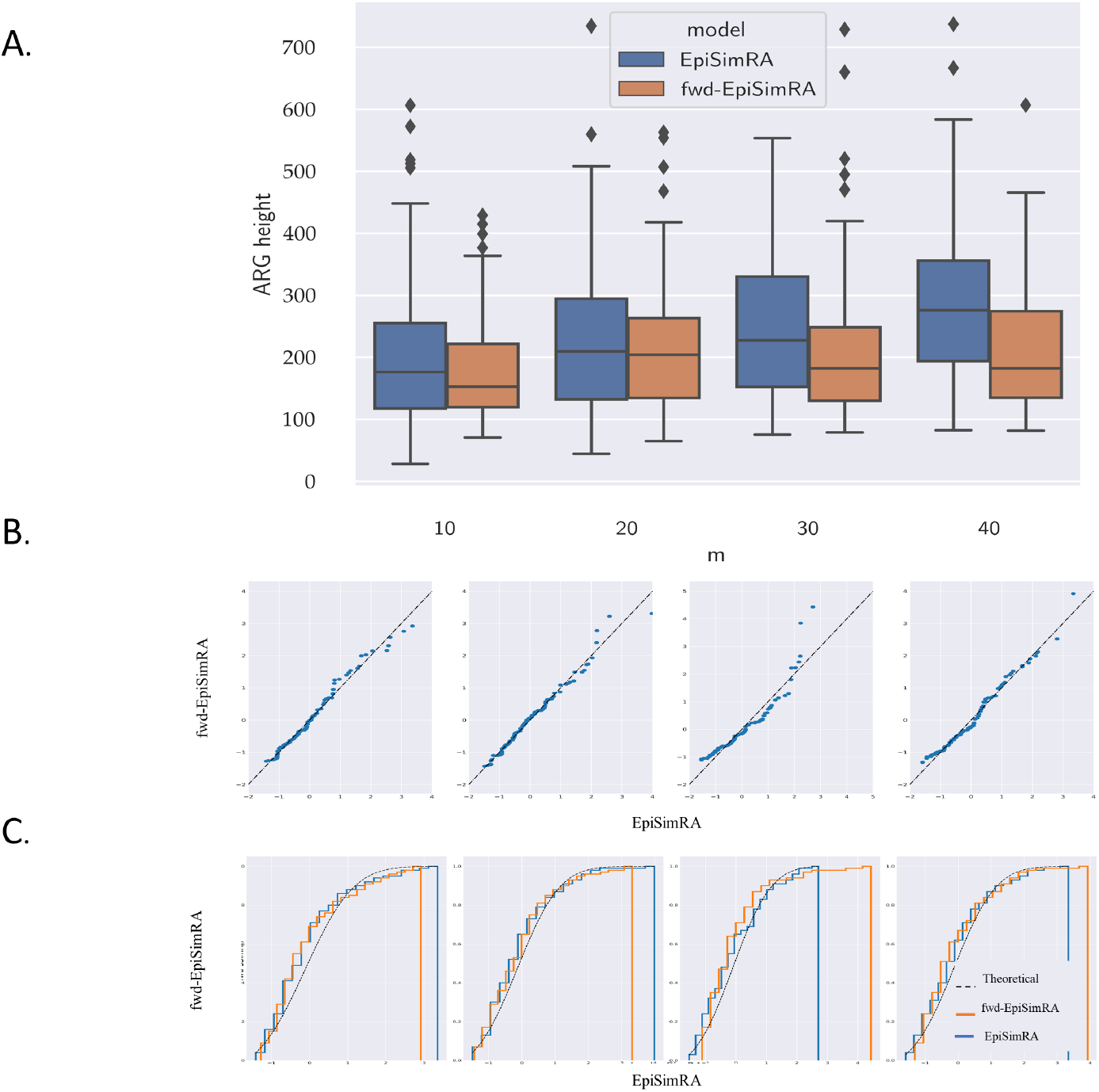
Comparison of the height of the ARG (H) between fwd-EpiSimRA and EpiSimRA with and without epistatis with recombination for *N* = 100, *g* = 250 kbp, *r* = 1.0 × 10^*−*8^, *m* = {10, 20, 30, 40}, *s* = {0.3, 0.3, 0.3} with epistastic parameters for *s*_*i*_*s*_*j*_ = 0.15 for *i, j* ∈ [1, 3] and *s*_1_*s*_2_*s*_3_ = 0.125. (A) The box-and-whisker diagram summarizes the result for each. On each box, the central mark is the mean, the edges of the box are the 25th and 75th percentiles, the whiskers extend to the most extreme data points not considered outliers, and outliers are plotted individually. (B) QQ plot and (C) Cumulative Distribution Function (CDF) plot of the backward and forward models show similar distributions with further agreement in Kolmogorov-Smirnov tests (Table B in Appendix).

#### Computational Complexity

Computational complexity of EpiSimRA is directly proportional to *N*, the number of individuals per generation; *g*, length of the genome under simulation and *k* for *k*-way epistatic interactions. As we increase these parameters, we obtain a more complex evolutionary history leading to longer running time due to longer time to coalescence for randomly sampled extant populations. As we observe concordance in the observed TMRCA for the coalescent simulator EpiSimRA as well as in its forward counterpart, fwd-EpiSimRA in the complex evolutionary scenarios when multiway epistasis is in effect, we obtain validation about the empirical correctness of the coalescent simulator. The coalescent simulator, EpiSimRA is extremely fast in finding approximations to TMRCA, in comparison to fwd-EpiSimRA, as the latter has to build the entire “book of populations” and trace it. This leads to a difference in running time of hours for the forward model vs. seconds for its coalescent counterpart with input dimensions from Table 1.

**Table 1.** Input parameters of the coalescent simulator

|  | Parameters | example values | user-specified units | units in bp for the algorithm | scaling factor |
| --- | --- | --- | --- | --- | --- |
| $g$ | segment length | 25; 75 | Kb | $\times 10^3$ bp | $\times 10^3$ |
| $m$ | extant units | 10; 20; 30; 40 | — | — | $\times 1$ |
| $N$ | population size | 100; 200; 500; 1000 | — | — | $\times 1$ |
| $I$ | length of genetic material | 1000 | bp | 1bp | $\times 1$ |
| rates/generation |  |  |  |  |  |
| $r$ | recombination rate | 1 | bp/gen $\times 10^{-7}$ | bp/gen | $\times 10^{-7}$ |
| $\mu$ | SNP mutation rate | 1.5 | mut/bp/gen $\times 10^{-8}$ | $\times 1$ mut/bp/gen | $\times 10^{-8}$ |
| selection, epistasis parameters |  |  |  |  |  |
| $s_i$ | fitness | 0.3 | — | $\times 1$ | — |
| $e_{ij}$ | epistasis | 0.1, 0.15 | — | — | $\times 1$ |

**Table 2.** K-S test statistics with corresponding p-values showing that the probability distributions of H as returned by *fwd-sSimRA* and *back-sSimRA* abstracts each other very closely.

| 3 interacting loci | | | $e_s$ | m | p-value | Test statistic |
| --- | --- | --- | --- | --- | --- | --- |
| $s_1$ | $s_2$ | $s_3$ | | | | |
| × | × | × | × | 10 | 0.1400 | 0.16 |
|  |  |  |  | 20 | 0.4431 | 0.12 |
|  |  |  |  | 30 | 0.3439 | 0.13 |
|  |  |  |  | 40 | 0.9995 | 0.05 |
| $s_1$ | × | × | × | 10 | 0.6766 | 0.08 |
|  |  |  |  | 20 | 0.7942 | 0.08 |
|  |  |  |  | 30 | 0.6766 | 0.10 |
|  |  |  |  | 40 | 0.5750 | 0.11 |
| $s_1$ | $s_2$ | × | × | 10 | 0.9921 | 0.06 |
|  |  |  |  | 20 | 0.5560 | 0.11 |
|  |  |  |  | 30 | 0.7942 | 0.09 |
|  |  |  |  | 40 | 0.8938 | 0.08 |
| $s_1$ | $s_2$ | × | 0.1 | 10 | 0.8938 | 0.08 |
|  |  |  |  | 20 | 0.9995 | 0.05 |
|  |  |  |  | 30 | 0.9710 | 0.06 |
|  |  |  |  | 40 | 0.7942 | 0.09 |
| $s_1$ | $s_2$ | $s_3$ | × | 10 | 0.3439 | 0.13 |
|  |  |  |  | 20 | 0.7942 | 0.08 |
|  |  |  |  | 30 | 0.6766 | 0.10 |
|  |  |  |  | 40 | 0.5576 | 0.11 |
| $s_1$ | $s_2$ | $s_3$ | 0.1 | 10 | 0.9610 | 0.07 |
|  |  |  |  | 20 | 0.9610 | 0.07 |
|  |  |  |  | 30 | 0.3556 | 0.13 |
|  |  |  |  | 40 | 0.6766 | 0.10 |

### 4.2 Evaluating Epistatic Scenarios

We compare the *H* under selection in EpiSimRA and show how different scenarios impact the height of the ARG (Figure 3 in Appendix). We find that positive selection affects the time to coalescence inversely with more selective pressure results in less time to coalescence when simulated with *N* = 1000 samples and genome length of *g* = 250 kbp. Epistasis in two and three interacting loci results in lower TMRCA than single locus selection and the neutral case amounting to higher selective pressure. We further studied effects of epistasis by simulating populations of *N* = 10, 000 with three-way epistasis. We find epistasis leads to a more complex evolutionary history resulting in longer time to coalescence in MRCA (Figure 4 in Appendix). When epistasis is not in effect i,e. when 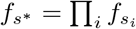, when *i* [1, 3] results in a shorter TMRCA with simpler evolutionary history. In addition, an exhaustive comparison between the two simulators for all scenarios with or without epistasis is included in the Appendix.

**Fig. 3.**
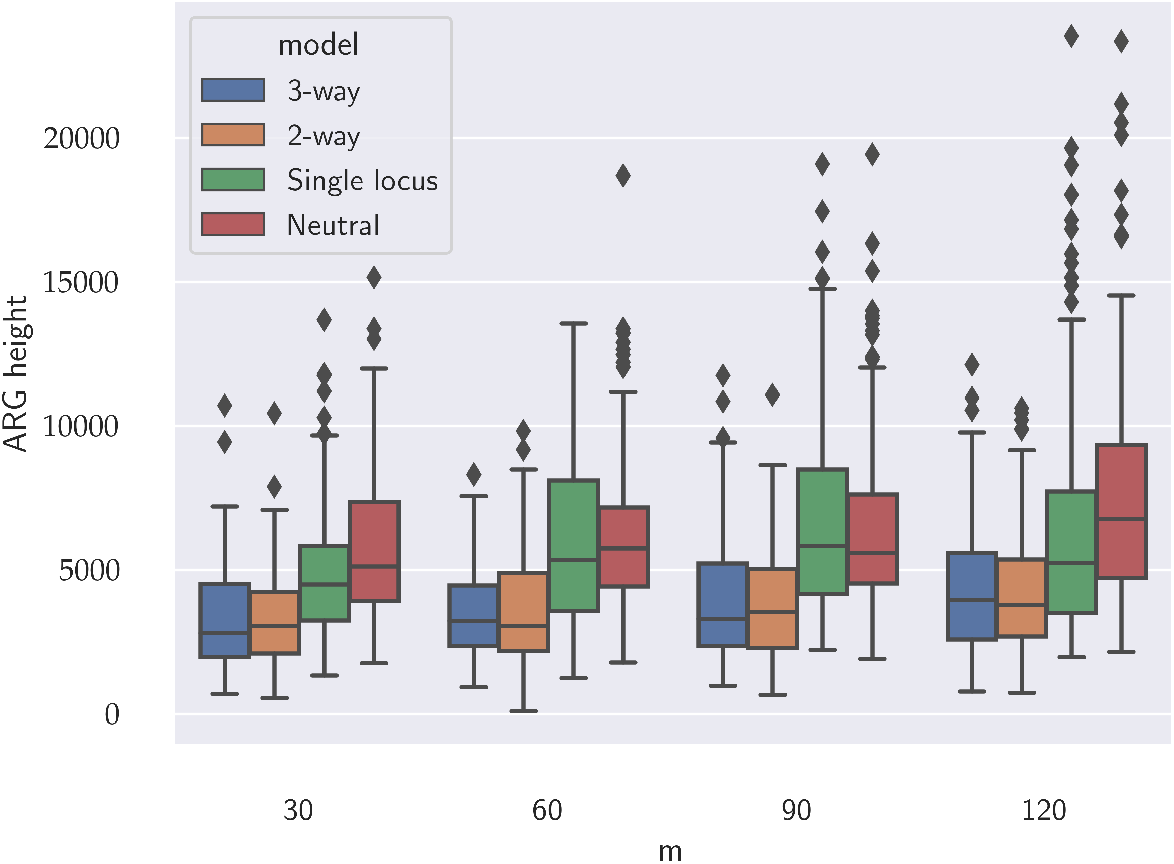
Comparing the height of the ARG (H) for different scenarios of selection in EpiSimRA with epistatis and recombination for *N* = 1000, *g* = 250 kbp, *r* = 1.0×10^*−*8^, *m* = {30, 60, 90, 120}, *s* = {0.3, 0.3, 0.3} with epistastic parameters for *s*_*i*_*s*_*j*_ = 0.15 for *i, j* ∈ [1, 3] and *s*_1_*s*_2_*s*_3_ = 0.125. The box-and-whisker diagram summarizes the result for each *m* and selection scenarios such as neutral (*s* = 0), single locus (*s* = 0.3), epistatic interaction at two loci and three loci respectively.

**Fig. 4.**
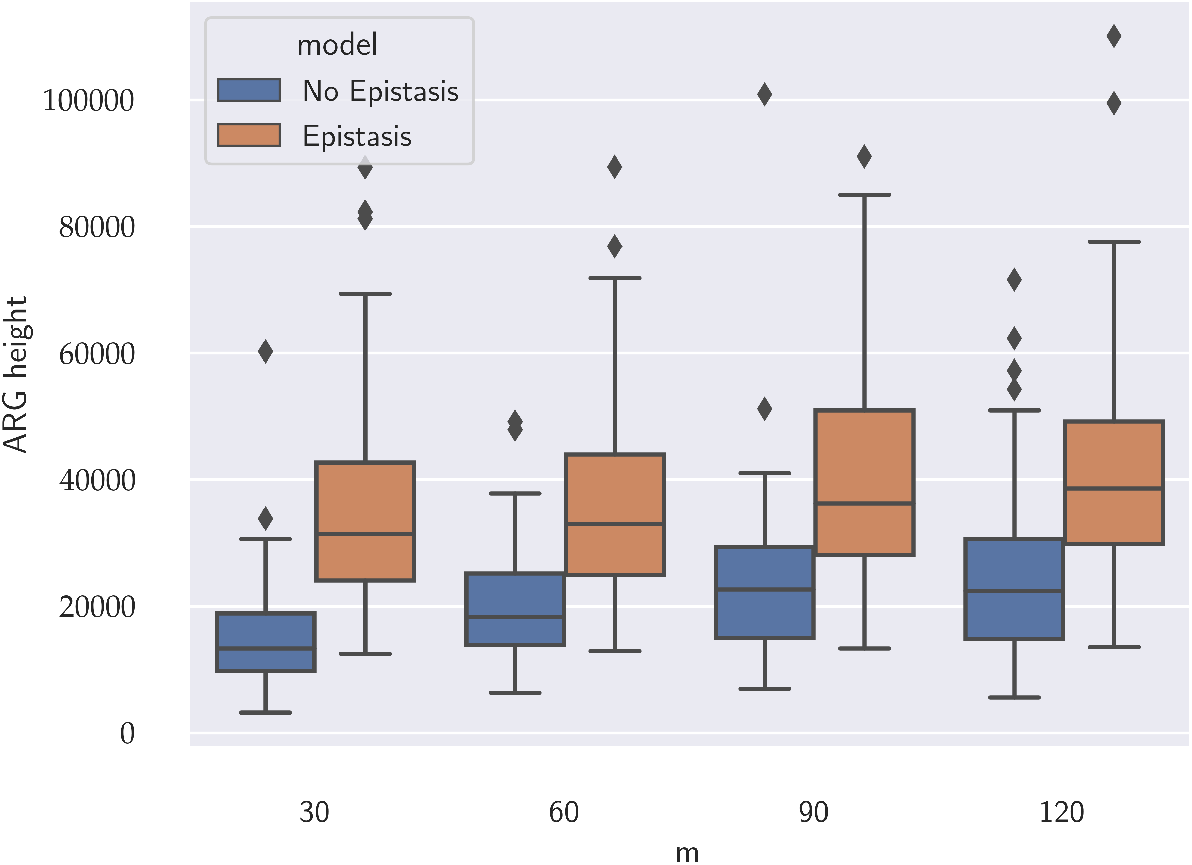
Comparing the height of the ARG (H) for different scenarios of selection in EpiSimRA with and without epistatis in recombination for *N* = 10, 000, *g* = 250 kbp, *r* = 1.0 × 10^*−*8^, *m* = {30, 60, 90, 120}, *s* = {0.3, 0.3, 0.3} with epistastic parameters for *s*_*i*_*s*_*j*_ = 0.15 for *i, j* ∈ [1, 3] and *s*_1_*s*_2_*s*_3_ = 0.125. The box-and-whisker diagram summarizes the result for each *m* and selection scenarios with and without epistatic interaction at three loci.

**Fig. 5.**
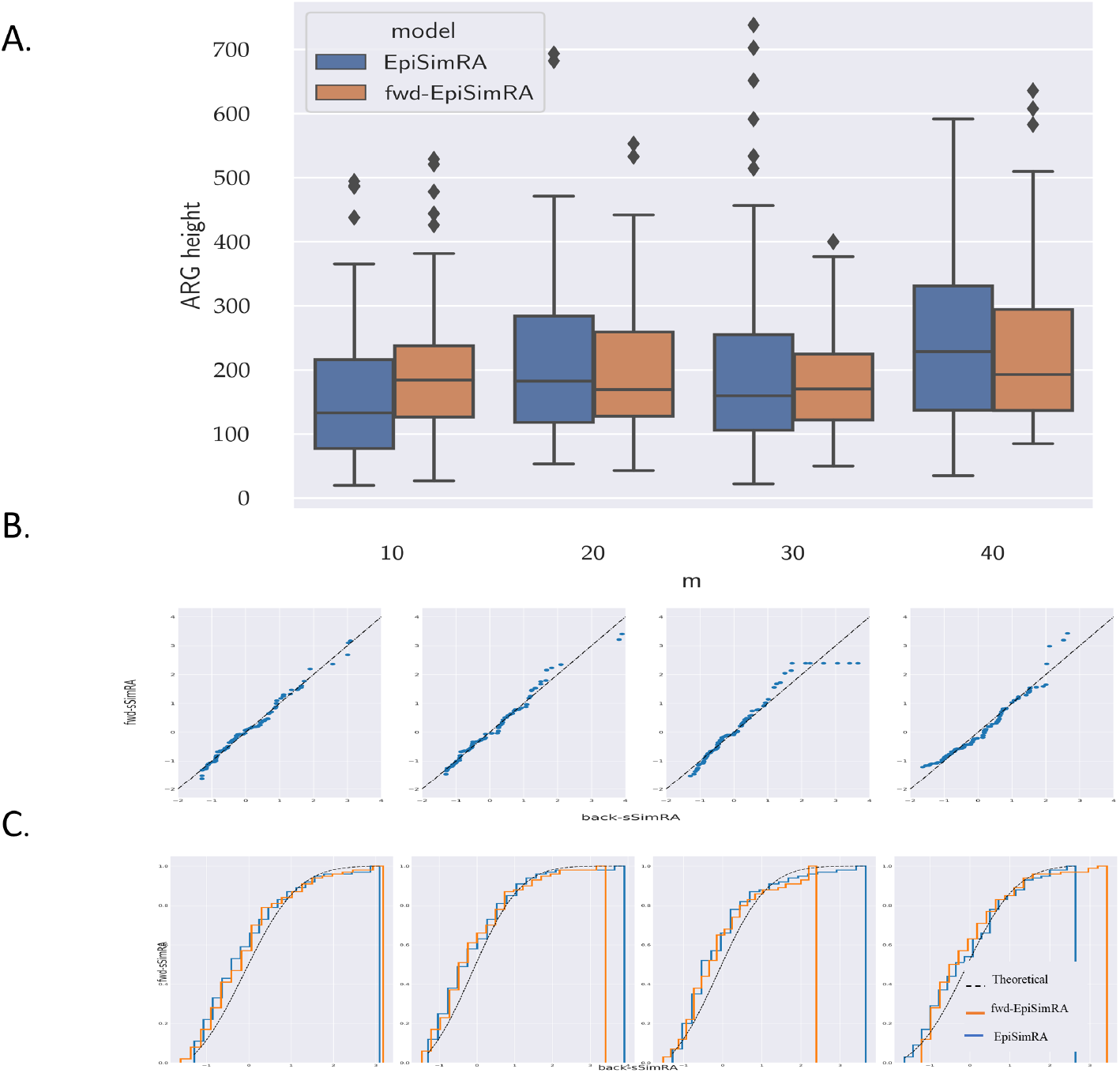
Comparing the height of the ARG (H) between fwd-EpiSimRA and EpiSimRA with and without epistatis in two loci with recombination for *N* = 100, *g* = 250 kbp, *r* = 1.0 × 10^*−*8^, *m* = {10.20, 30, 40}, *s* = {0.3, 0.3} with epistastic parameters for *s*_0_*s*_1_ = 0.15. (A) The box-and-whisker diagram summarizes the result for each. On each box, the central mark is the mean, the edges of the box are the 25th and 75th percentiles, the whiskers extend to the most extreme data points not considered outliers, and outliers are plotted individually. (B) QQ plot and (C) CDF plot of the backward and forward models show similar distributions.

**Fig. 6.**
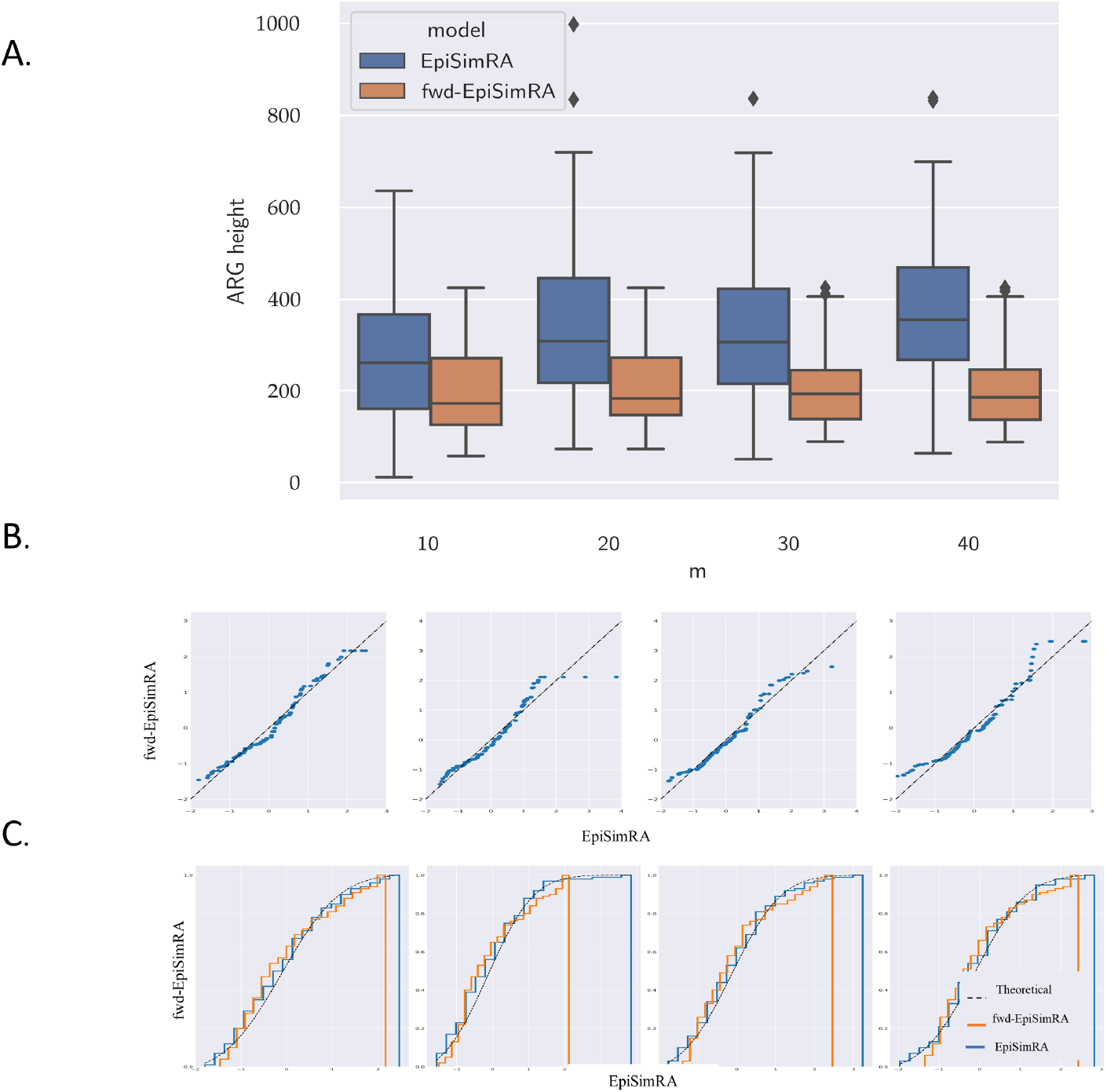
Comparing the height of the ARG (H) between fwd-EpiSimRA and EpiSimRA for selection in single locus with recombination for *N* = 100, *g* = 250 kbp, *r* = 1.0 × 10^*−*8^, *m* = {10.20, 30, 40}, *s* = 0.3. (A) The box-and-whisker diagram summarizes the result for each. On each box, the central mark is the mean, the edges of the box are the 25th and 75th percentiles, the whiskers extend to the most extreme data points not considered outliers, and outliers are plotted individually. (B) QQ plot and (C) CDF plot of the backward and forward models show similar distributions.

**Fig. 7.**
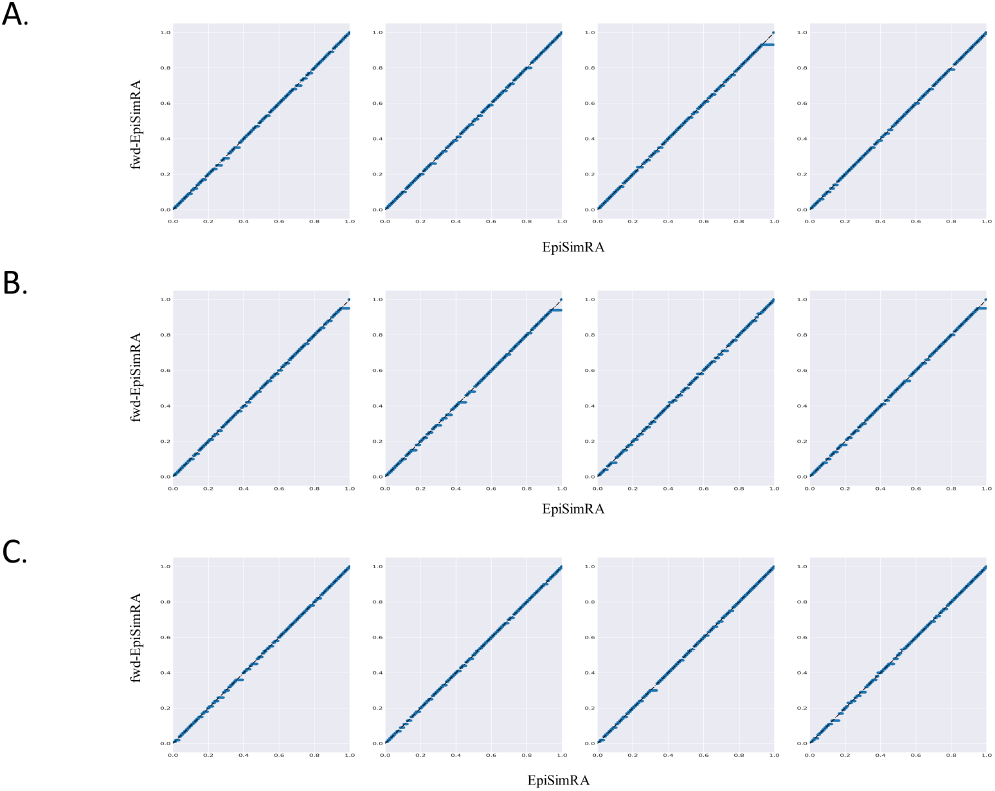
P-P plots of distributions of the height of the ARG (H) between fwd-EpiSimRa and EpiSimRA for (A) single locus selection, (B) epistatic interaction at two loci and (C) epistatic interaction at three loci *g* = 250*K, r* = 1.0 × 10^*−*8^ *N* = 100, *s* = 0.3, *e*_*s*_ = {0, 0.1} and *m* = {10, 20, 30, 40}.

## 5 Conclusions

We present an algorithm that builds multi-locus selection and multiway epistasis into the backward coalescent model with recombinations, as well as, in a forward scheme. Moreover, to the best of our knowledge, this is the first model which took a backward simulator with multiway epistasis and compared it nose-to-nose with its forward counterpart. Through extensive empirical comparison studies, albeit for small populations due to the time constraint of the forward model, we show that for complex scenarios with selection and epistasis (or even under neutral scenarios) the hallmark values by the backward and the forward schemes approximately abstract each other. As the distributions of both the schemes are concordant, we conclude that any one of the simulators (EpiSimRA or fwd-EpiSimRA) can be used to understand the effects of negative and positive selection, with multiway epistasis, along with selective sweeps across generations. Due to lack of similar assumptions, parameters and hallmarks of ARGs returned we did not compare EpiSimRA with present coalescent simulators for selection at a single locus. As fwd-EpiSimRA closely imitates Wright-Fisher model and allows for epistatic interactions, we used it as a validation framework.

Multiway interaction across multiple loci leads to complex population genetic history with a longer height of the ARG relative to non-epistatic interactions. EpiSimRA encompasses all such scenarios with the potential for further exploration for viral phylodynamics with randomly sampling a bacteria or virus populations. The time to coalescence when reconstructing its phylogeny under selection and epistasis allows us to study important epidemiological, immunological and evolutionary processes of viruses [30] such as the recent SARS-CoV-2 or similar Coronaviridae. This allows a validation framework for including selection and epistasis into standard population genetic models where we can now study the different scenarios when all the diploids associated with mutated sites along the chromosome with differing fitness values corresponding to the alleles.

## Appendix

### A The Forward Simulator

For two individuals *i* and *i*′, the ratio of the probabilities that a locus *l* contributes to whether an individual will have an offspring is the relative fitness

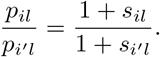

The total ratio of probability *i* will have children to *i*′ having children is

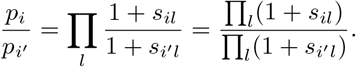

From this, it follows that

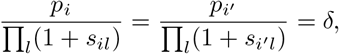

where *δ* has the same for all *i, i*′ and all other individuals.

Given this,

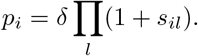

SinceΣ _*i*_*p*_*i*_ = 1 = *δ Σ*_*i*_ *П*_*l*_(1 + *s*_*il*_), it follows that

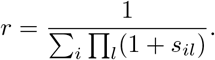

Therefore, the probability that *i* has children is

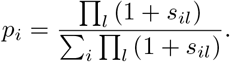

For multiway epistasis we include the conditions of *k* loci being linked with each other such that a combined fitness coefficient is calculated by taking the fitness product at each interacting site as defined in Equation 8.

#### Choosing Parents

The probability that two children will pick the same parent is operationally, the reciprocal of the effective population size [7,17]. Likewise, the same interpretation was made by [19] in the construction of the coalescent. Given a set of *p*_*i*_’s in a given generation *t*, the probability that two children will pick the same parent is 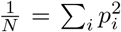. While the *p*_*i*_’s define the probability that children pick their parents, *N* does not play a direct role in determining the course of the algorithm in constructing the book of populations but will affect the shape of the ARG that is traced in the second stage.

With selection on a single locus in effect, each generation will have *N*_*s*_ individuals that contains the allele under selection, yielding Π_*l*_(1 + *s*_*il*_) = 1 + *s*, and (*N* − *N*_*s*_) individuals without the allele with Π_*l*_(1 + *s*_*il*_) = 1.

#### Transfer of genetic material

After the children have randomly selected their parents, the child requests one chromosome from each of the parents. The parents randomly select whether to pass one of their two chromosomes, or to construct a new chromosome via a recombination event involving a crossover between it’s two chromosomes with respect to the recombination rate, *r*. If a crossover is generated, the parent randomly selects a location and transfers the genetic material upto that location from one chromosome and the rest from the it’s other copy. This is done in part to reconstruct the ARG, and to characterize genetic variation along chromosomes yielding the final recombinations [27]. In case of no recombination, the parent randomly decides which chromosome’s genetic material should be passed over to the child (see Fig. 1 of the main manuscript).

Each newly constructed chromosome is painted with new SNP mutations randomly generated according to a mutation rate probability *µ*, a randomly selected location, and allele value. With probability of mutation on each polymorphic site, the resultant mutated chromosomes are finally passed to the child from the parents along with the sites of mutations and recombinations.

Throughout the generation, forward in time, we keep track of the sites of recombinations and mutations to efficiently trace the ARG from extant individuals to it’s GMRCA.

#### Tracing the ARG from the book of populations

Detecting the past recombination events from extant sequences and specifying the place of each recombination and recombinant sequences has been well studied [14,21,24]. The ARGs define a genealogical graph for all of the chromosomes in a population. For each locus, the ARG for any given segment between recombination crossovers will form a tree. When the sequences are non-recombining, we only need to use coalescences and mutations to describe their genealogy to find a most recent common ancestor (MRCA). Traversing back through an ARG, coalescent events are very common in occurrence, but, in case of a recombination, the history of lineages not only show bifurcations, but also recombinations resulting in cycles. Our algorithm looks for recombination events going back every generation and traces them until convergence to a GMRCA.

## A.1 Simulating the book of populations with selection and two-way epistasis

### ALGORITHM

1. **Initialization**
  a. *N* individuals (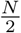 males and 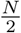 females) in the base generation, which remains constant throughout the simulation.
  b. Number of Generations, *G* = *c* \**N*, where *c* is a constant.
  c. Randomly allocate genetic material along the length of chromosome, *g*.
  d. Assign selection coefficients for interacting sites for two-way epistasis (0 for neutral).
  e. Set flag, *f*, for allele(s) under selection on a mutated site (0 for neutral).
2. If *f* is set, randomly select an individual among *N* and a site, *g*_*s*_ along *g* which underwent mutation. Select an allele randomly in *g*_*s*_ and set *f* to 1.
3. **Loop** For each generation, *t* ∈ {1, …, *G*}
4. **Loop** For each individual *i* in {1, …, *N*}, in (*t* − 1)^*th*^ generation.
5. Compute 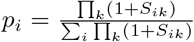, where any group *k* of loci could contain a single locus under selection, for which *S* = *s* is defined as the user input. It can also contain a locus interacting with another locus, in a two-way epistasis. In this case *s* is populated from a matrix formed by the all possible alleles at each loci, from the following form, 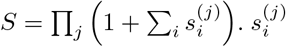 is the selection cofficient at allele *j* in individual *i*’s chromosome.
6. Select parents for each child in *t*^*th*^ generation based on *p*_*i*_ from (*t–* 1)^*th*^ generation.
7. **End**
8. For each child *i* in *t*^*th*^ generation, compute scaled recombination rate *r*^*t*^ = *r* * *g* and select a value, *r*_*val*_ ∈ [0, 1].
9. If 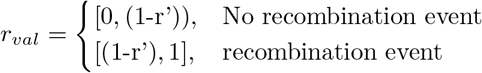
10. If No recombination event: Randomly pick a chromosome from the parent and assign it’s genetic material to the child.
11. Else Randomly pick a crossover index *z*∈[1, *g*]. Get the genetic material from [1, *z*] in the first chromosome of the parent and [(*z* +1), *g*] in the second, combine them and assign it to the child.
12. In the child’s genetic material, randomly select locations along the chromosome length, *g* for mutation according to the Poisson distribution and the scaled mutation rate *µ*′ = *µ*\**g*. Assign the alleles randomly to other bases. For example, if the allele was *A*, change it randomly to one of the other bases {*G, T, C*}.
13. Update the Chromosomes of the current generation with the new genetic information obtained from the previous generation and continue until the last generation, *G*.
14. **End**

## A.2 Tracing the ARG from the book of populations

### ALGORITHM

1. **Initialization**:
  a. Randomly select *m* number of extant individuals from *N* in the last generation.
  b. Select one chromosome out of the two in these *m* extant samples, randomly. Compute the active lineages, *j* by comparing the genetic material *g* in each of the *m* chromosomes selected.
2. **Loop** for each generation, *t* going backwards from {*G*, …, 1}
3. Identify each chromosome from the previous generation (*t*−1) which contributed to each chromosome in the current generation, following the book of populations.
4. Check to see if multiple children in the *g*^*th*^ generation share the same parent in the previous generation.
5. Iterate and Count the number of active samples, *m*′ in each generation.
6. **Until** *m*′ = 1
7. Compute the Height of the GMRCA from the height of convergence.

### B Experiments and Comparison study

Here we exhaustively list all the box-whisker diagrams, Q-Q and CDF plots for two-way epistasis and P-P plots for all experiments conducted while comparing the two simulators fwd-EpiSimRA and EpiSimRA (Figure B5).

We also provide the test statistics and p-values obtained by running K-S test which does not reject the null hypothesis that the samples of *H* as returned by the two simulators are indeed drawn from the same distribution as shown in Table B.

